# Altered parasitism of a butterfly assemblage associated with a range-expanding species

**DOI:** 10.1101/2020.02.13.947440

**Authors:** H. Audusseau, N. Ryrholm, C. Stefanescu, S. Tharel, C. Jansson, L. Champeaux, M. R. Shaw, C. Raper, O. T. Lewis, N. Janz, R. Schmucki

## Abstract

**Aim:** Biotic interactions are an important factor structuring ecological communities but data scarcity limits our understanding of the impact of their response to climate and land use changes on communities. We studied the impact of a change in species assemblage on biotic interactions in a community of closely-related butterflies. Specifically, we examined the impact of the recent range expansion of *Araschnia levana* on the resident species, with a particular focus on natural enemies, parasitoids, shared with other butterfly species in the assemblage.

**Location:** Sweden.

**Time period:** Two years (2017-2018).

**Major taxa studied:** Nettle-feeding butterflies (*Aglais urticae, Aglais io, Araschnia levana*, and *Vanessa atalanta*) and their parasitoids.

**Methods:** We assessed parasitism in 6777 butterfly larvae sampled in the field from 19 sites distributed along a 500 km latitudinal gradient, and every two weeks throughout species’ reproductive seasons. We identified the parasitoid complex of each butterfly species and their overlap, and analysed how parasitism rates were affected by species assemblage, variations in abundance, time, and the arrival of *A. levana*.

**Results:** Parasitoids caused high mortality, with substantial overlap across the four host species. The composition of the host community influenced parasitism rates and this effect was specific to each species. In particular, the rate of parasitism in resident species was comparatively higher at sites where *A. levana* has been established for longer.

**Main conclusions:** Parasitoid pressure is a significant source of mortality in the nettle-feeding butterfly community studied. Variations in butterfly species assemblages are associated with substantial variations in rates of parasitism. This is likely to affect the population dynamics of their butterfly host species, and, potentially, the larger number of species with which they interact.

## Introduction

Biotic interactions are important drivers structuring ecological communities. While occurring locally, the impact of biotic interactions is visible across ecological scales, influencing population dynamics, determining community structures and patterns of species co-occurrence, and shaping distribution ranges and abundances (Araújo & Luoto, 2007; Heikkinen et al., 2007; Meier et al., 2010; Wisz et al., 2013; Belmaker et al., 2015). Biotic interactions can nevertheless be altered by climate and land use changes, thereby disrupting ecological communities (Tylianakis et al., 2008; Blois et al., 2013). The importance of biotic interactions is widely recognised in the literature, including its importance for refining predictions of species’ responses to environmental change (Wisz et al., 2013; Dormann et al., 2018), but the scarcity of comprehensive empirical data strongly limits our ability to understand the larger-scale impact of their response to environmental change on populations and communities. Most studies that examined changes in ecological networks along environmental gradients are based on correlative approaches (Pellissier et al., 2017) and, therefore, cannot disentangle the effect of biotic interactions from the effects of environmental change. Thus, measuring and understanding the impact of biotic interactions in a context of change remains a major challenge.

This difficulty is partly due to the dynamic nature of biotic interactions and the many ways and different scales that environmental change can affect biotic interactions (Wisz et al., 2013; Kissling & Schleuning, 2015; Pellissier et al., 2017; Dormann et al., 2018). For example, changes in climate and land use can affect the distribution and demography of species, which in turn might alter the nature and strength of biotic interactions and their impact on ecological communities and species distribution (Tylianakis et al., 2008; Early & Keith 2019). Differences in sensitivity of interacting species to environmental changes can 1) alter their respective abundances and/or spatial pattern of co-occurrence, 2) disrupt the temporal synchrony between them, or 3) induce new interactions in the case of the establishment of invasive or alien species.

Understanding and identifying the forces that shape biotic interactions in changing environments is particularly important for invertebrates such as insects, a group that constitutes more than half of the biodiversity of Earth and underlies ecosystem services that directly contribute to ecosystem productivity and stability, and to human well-being (see Losey & Vaughan, 2006). While direct interactions such as predator-prey and insect-plant interactions are widely studied and documented, the vast majority of interactions operate in complex networks where species are connected through both direct and indirect interactions. The process of apparent competition is an example of an indirect interaction, where the population dynamics of species at the same trophic level can be linked via the action of shared natural enemies (Holt & Lawton, 1993; 1994). For example, the invasion and establishment of a closely-related insect species can be detrimental to a native species by increasing the resources available for their shared parasitoids. Apparent competition mediated by shared parasitoids was shown for leafhopper communities in California where the introduction of a new host species, combined with the strong preference for the native species, resulted in an overall increase in parasitoid pressure and decline of the native species (Settle & Wilson, 1990). In contrast to direct interactions, indirect interactions are generally more complex and involve several trophic levels, which makes their identification, as well as the evaluation of their effects on species and communities, more difficult. Laboratory experiments conducted on *Drosophila* assemblages in microcosms have shown that changes in biotic interactions (both direct and indirect) along a climatic cline influence population dynamics (Davis et al., 1998). In natural systems, the impact of apparent competition has been shown to vary with the size of the community, the abundance of hosts and their phenology (Bonsall & Hassell, 1997; Van Nouhuys & Hanski, 2000; Morris et al., 2004; Blitzer & Welter, 2011), and can affect multiple species that share common enemies (Morris et al., 2004, Frost et al., 2016). However, our understanding of indirect interactions is mainly derived from a small amount of experimental data gathered under laboratory conditions or at relatively small spatial and temporal scales. The lack of detailed data collected across regions and over multiple generations limits our ability to quantify and predict the impacts of indirect biotic interactions on populations and communities in the context of environmental change.

Here we focus on apparent competition mediated by shared parasitoids in a community of closely-related (Nymphalidae: Nymphalinae, Nymphalini) nettle-feeding butterflies (*Aglais urticae*, *Aglais io*, *Vanessa atalanta*) along a latitudinal gradient in Sweden. We investigate the impact of the range expansion of *Araschnia levana*, a newly-arrived butterfly also feeding on nettle (*Urtica dioica*). The establishment and expansion of *A. levana* in Sweden is most likely a result of the warmer conditions observed over the last decades and has the potential to modify the interactions that structure the community of resident nettle-feeding butterflies. Recent analyses of species co-occurrence of three butterfly species (*Aglais urticae, Aglais io*, and *A. levana*, Audusseau et al., 2017) have shown the potential effect of the newly-established species in southern Sweden on the distribution and niche partitioning of the resident species. Audusseau et al. (2017) observed a shift in the distribution of *A. urticae* and *A. io* following the establishment of *A. levana* in Sweden, and suggested that these shifts could be explained by apparent competition, mediated by shared parasitoids. To further investigate this hypothesis and document the impact of such a change in species assemblage, we conducted a field study spanning a 500 km latitudinal gradient in Sweden, along the establishment gradient of *A. levana*. We investigated the phenology of parasitism of the nettle-feeding butterflies and its spatio-temporal structuring, and whether the change in parasitism rate was linked to a change in potential for apparent competition that the resident species experienced when co-occurring with the newly-established species.

## Material and Methods

### Study system

*Aglais urticae, Aglais io, Araschnia levana* and *Vanessa atalanta* are closely-related butterfly species from the same tribe (Nymphalini) within Nymphalidae family. The larvae of all four species feed (practically exclusively) on nettle (*Urtica dioica*), but they differ in their egg-laying behaviour, phenology, and distribution.

*Aglais urticae, A. io*, and *A. levana* are batch-laying species, with batches of 10 to 40 eggs for *A. levana* and of 200-300 eggs for *A. urticae* and *A. io*, while *V. atalanta* lays eggs singly (Ebert 1993). *Aglais urticae* and *A. io* batches are laid at the apex of nettle plants. During the first three instars of their development, the larvae are gregarious and conspicuous as they feed near the apex of the nettle stem. At their fourth instar, larvae become solitary and feed over all of the plant and may hide in the foliage. Larvae of *A. levana* are also gregarious in the early instars and become solitary later on. However, batches of this species are less conspicuous to the human eye. The smaller size of both the batches and the larvae causes less damage to the plant, and larvae often feed on the lower surface of the leaf.

The four species have broadly overlapping phenologies, with adults flying from March to September. However, due to differences in voltinism (the number of generations a species has every year) and yearly variations in weather conditions, the time periods during which larvae of each species are found in the field vary. Populations of *A. urticae* are bivoltine in south Sweden and becomes progressively univoltine as we move to the northernmost part of the country. Larvae of this species are recorded from early May to the end of August. Butterfly individuals from the first generation correspond to the eggs laid in May. In the south, individuals from the second generation are the offspring of adult butterflies from the first generation and correspond to eggs that start to be laid about six weeks later. In between these two peaks, the abundance of *A. urticae* larvae drops. *Aglais io* is univoltine in Sweden and starts reproducing soon after *A. urticae*, with larvae observed from late May to early August.

*Araschnia levana* is an obligate bivoltine species. In contrast to *A. urticae* and *A. io*, which overwinter as adults, individuals of *A. levana* hibernate in the pupal stage. Butterfly larvae from the first generation are found in the field in June; larvae from the second generation are found from end of July to early September. Last, *V. atalanta* is a migratory butterfly in Sweden and its population depends on the migratory influx from the areas where the species is resident. It is univoltine in Sweden with larvae observed in the field from May to early September.

These species are distributed over most of Sweden, except for *A. levana* whose distribution is so far limited to the southern half of the country. The first anecdotal observations of *A. levana* were reported in the county of Skåne in Sweden in 1982 and the species is now well-established in the southern part of the country (Eliasson et al., 2005). Further, opportunistic occurrence data extracted from Artportalen (Swedish Species Observations System, www.artportalen.se, 30/08/2019) showed that the species has progressively expanded from the county of Skåne to Kronoberg and further north, but has not yet reached the Stockholm area (most northely latitude of observation in 2017: 58.6981, see Appendix S1 in Supporting Information).

### Field sampling

We collected larvae of the four study butterflies, *A. urticae, A. io, A. levana*, and *V. atalanta*, over two years (2017-2018) and fortnightly throughout the species’ reproductive season (May-August). Our sampling was distributed across 19 sites along a 500 km latitudinal gradient from south Sweden to the Stockholm area (Fig. S1). The 13 sites located in the southern part of Sweden fall within the distribution range of all four butterfly species, while the six sites in the Stockholm area are north of *A. levana’*s current range.

### Larval sampling and monitoring

We focused on larval parasitism. Pupal parasitism is also likely to cause high mortality in the species studied (Pyörnilä, 1977; Shaw et al., 2009), but the solitary and concealed pupae are difficult to collect in sufficient numbers for reliable estimates of pupal mortality. To maximize the diversity of captured parasitoid species, we sampled butterfly larvae at different developmental stages. We followed such a stratification of the sampling effort because the temporal window of attack of butterfly larvae differ among parasitoids species and can be restricted to a few developmental stages. For example, while ichneumonids of the genus *Thyrateles* attack very late larval or prepupal stages, *Cotesia vestalis*, which can be an important opportunistic parasitoid of at least *A. urticae*, parasitizes first instar larvae and emerges mainly from second instar larvae (MRS, personal observation). Therefore, at each sampling occasion and for each butterfly species we aimed to collect seven second instar larvae per batch from a maximum of five batches, 20 fourth instar larvae per batch from a maximum of five batches, and up to 20 fifth instar larvae, where possible from different batches.

We kept the collected larvae in transparent plastic boxes (155×105×45mm) with up to five individuals from the same batch in a box. We reared larvae under laboratory conditions (temperature 23°C, light regime 22L:2D) and fed them daily with *Urtica* leaves from location from where the larvae had been collected. This was because some of the Tachinidae parasitoids (*Sturmia bella* and *Pales pavida*, of those encountered) lay microtype eggs on nettle leaves and the butterfly larvae become parasitized only when they eat the infected leaves.

For larvae that were parasitized, we recorded the date and stage from which the parasitoid emerged (larval instar or pupa). We kept parasitoids individually or per batch in plastic vials, under the same laboratory conditions as the butterfly larvae. We preserved freshly-dead adult parasitoids in 95% alcohol, before taxonomic identification. The parasitoid pupae that did not hatch by early September, as well as the pupae from the second generation of *A. levana* (which have an obligate diapause before adult emergence), were kept cool during the winter period, until we broke their diapause around mid-April (see Appendix S2 for details on the diapause conditions).

### Analyses

We performed all analyses in R 3.6.1.

#### Parasitism rates across counties

We investigated variation in overall parasitism rates per butterfly species and county (Skåne, Kronoberg, and Stockholm). We performed this analysis in a Bayesian framework, using generalized linear and nonlinear multivariate multilevel models. We modelled parasitism rate assuming a binomial distribution and a logit link function. We tested for the effect of species, county, year, and the interaction between species and county as linear effects on parasitism rate and included the week of sampling as a non-linear effect (with k up to 4) to control for phenological variations in parasitism rate for each species. We grouped sites by county (Skåne, Kronoberg, and Stockhom, see Fig. 1) to reflect the south-north progression of the establishment of *A. levana*, and increase the power of our analyses along this gradient. We fitted the model through MCMC sampling, using the Hamiltonian Monte Carlo algorithm implemented in Stan (Carpenter et al., 2017) and the R interface provided in the brms package (Bürkner 2017; 2018). We ran four chains for 10000 iterations with the first 4000 discarded as burn-in and used the default non-informative priors. To test for significant differences in parasitism between county and species, we compared the posterior probability distribution of the model parameters.

**Figure 1.**
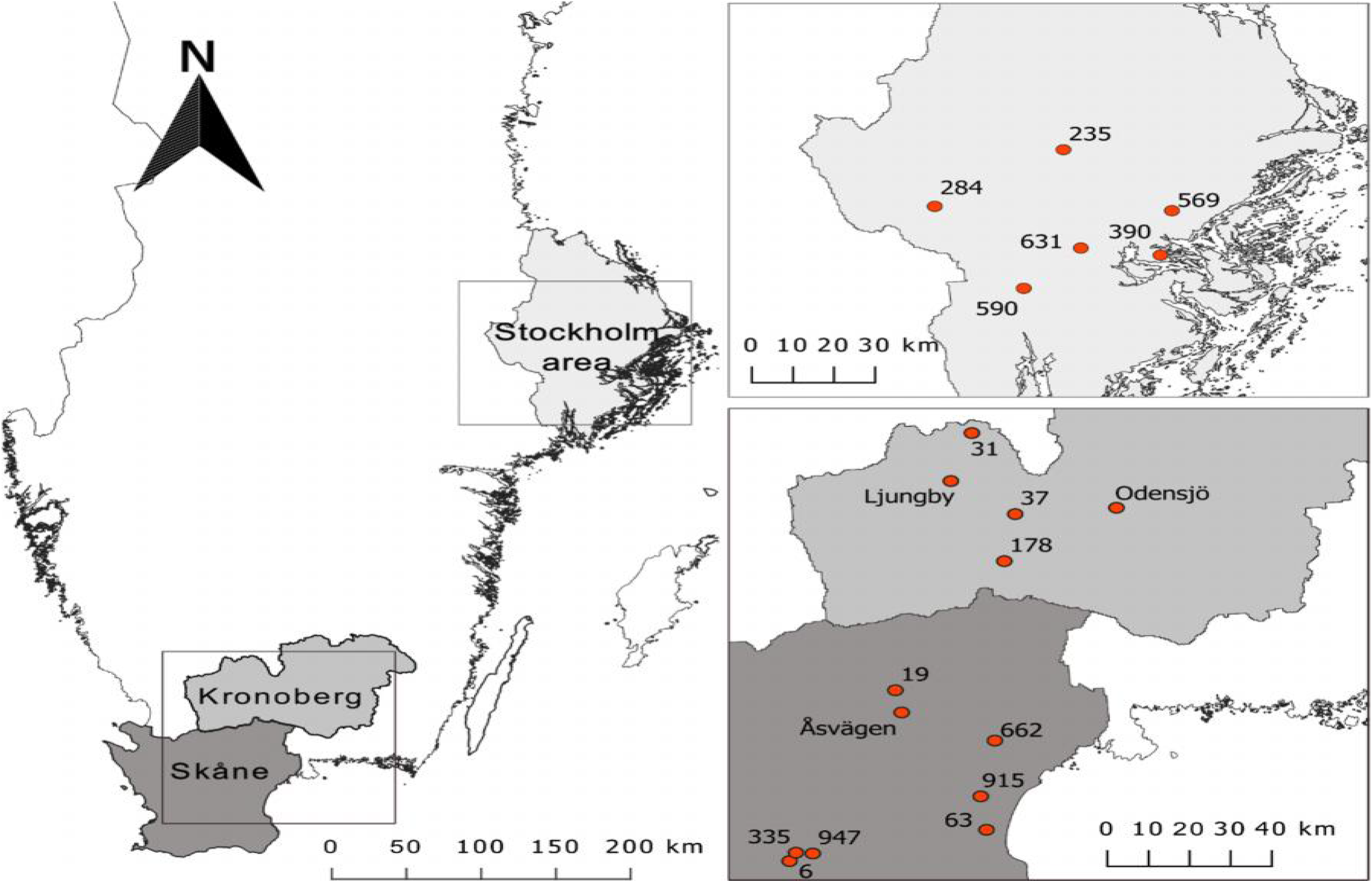
Map showing the 19 sites, spread across three counties, visited fortnightly over the two field campaigns (2017-2018).

#### Butterfly community and parasitism rate

We examined the effect of the butterfly community composition on each species’ parasitism rates. Specifically, we tested for the effect of the presence or absence of each species of butterfly, taken as a binary variable (0/1), and the effect of the abundance of larvae, on the parasitism rate of focal species. We also included in each model the non-linear effect of the sampling week (with k up to 4), to capture phenological variations of parasitism of each species. The abundance of larvae corresponds to the total number of larvae from all species collected per site and sampling week and was zero-centred prior to inclusion in the models. We performed these analyses in a Bayesian framework, using generalized linear and nonlinear multivariate multilevel models. Parasitism was modelled assuming a zero inflated binomial distribution with a logit link function and we used the same parameters as previously mentioned for model fitting. Lack of data on parasitism of *A. levana* prevented us from investigating the impact of community composition on parasitism for this species. Note that this analysis examined the effect of the butterfly community composition on each species’ parasitism rates, regardless of the parasitoids responsible for the parasitism rate recorded. Since the parasitoids responsible for the highest mortality are partially or entirely shared between the study butterflies (Table 2), these analyses explored how each species co-occurrence is linked to the parasitism of each of the focal species, by the likely action of parasitoids. These analyses do not investigate, however, the specific impact of the arrival of *A. levana* on the parasitism of native species. In particular, because if apparent competition participates to the structuring of this butterfly community, it can only occur via the action of natural enemies shared between the native species and *A. levana*. In Appendix S3, we offer a specific analysis of the potential impact of *A. levana* on parasitism of native species, examining the effect of the butterfly community composition on parasitism rates by the subset of parasitoids shared with *A. levana*.

#### Parasitism rate and time since establishment of A. levana

We investigated the role of the establishment of *A. levana* on parasitism rate of the native species. The available observations of *A. levana* clearly suggest that the species has first established in the southern part of the country and is progressively moving northward (see Appendix S1). If the establishment of *A. levana* has induced an increase in parasitism rate in the native species through apparent competition (as proposed by Audusseau et al., 2017), we would expect a decrease in parasitism rates with latitude. In addition, as the establishment of *A. levana* and its progression might not strictly follow the latitudinal gradient and could aslo be influenced by the configuration of landscape features such as presence of corridors or barriers affecting their dispersal, we also tested for the effect on parasitism rate of the time since first observation of *A. levana* within a 10 km buffer zone around each site and hypothesized that there would be a negative correlation between parasitism rate and the time since first observation.

For each species, we tested the effects of latitude and time since the first observation of *A. levana* in the 10 km the buffer zone around the sites, using generalized linear models and assuming a binomial distribution. We restricted these analyses to sites where *A. levana* is now established (Skåne and Kronoberg). Data on time since colonization were extracted from Artportalen (Swedish Species Observations System, http://www.artportalen.se/, 30/08/2019). The latitude and the time since first observation of *A. levana* at a site are closely correlated, as they both reflect the south-north gradient of progression of *A. levana*. Therefore, we transformed the time elapsed since the first observation of *A. levana* into a 4-level ordinal variable, which corresponds to the division of the distribution of this variable into 4 quartiles, to group sites by periods of establishment of the invading species. The dates of the first, second and third quartiles were 16/05/2004, 22/07/2006 and 02/08/2007. Latitude was zero-centred before it was included in the model.

## Results

### General patterns of incidence of butterfly species and parasitoid attack

Over the two sampling seasons, we sampled 6777 butterfly larvae across the 19 sites (*A. io* = 2259, *A.urticae* = 2254, *A. levana* = 1583, *V. atalanta* = 681). The three most widespread butterfly species occurred at all sites, except *A. io* which was absent at three sites (Odensjö, Åsvägen, 31), and *A. urticae* which was absent from site 31. As expected, *A. levana* was not observed at the latitude of the Stockholm area but it was found at all sites further south.

Of the 6777 collected larvae, 1508 were parasitized and produced parasitoids from three families: Tachinidae (Diptera), Ichneumonidae (Hymenoptera) and Braconidae (Hymenoptera). We identified 11 species: the tachinids *Pelatachina tibialis, Sturmia bella, Phryxe vulgaris, Phryxe nemea, Pales pavida* and *Blondelia nigripes*, the ichneumonids *Phobocampe confusa, Thyrateles haereticus* and *Thyrateles camelinus*, and the braconids *Microgaster subcompleta* and *Cotesia vanessae* (Table 1, 2). Despite collecting very early instar larvae, we did not encounter *Cotesia vestalis*. Overall, 76.7% of the parasitized larvae were parasitized by either *P. tibialis, P. confusa* or *S. bella*, which represented 34.6, 28.5 and 13.6% of the cases of parasitism, respectively. *Pelatachina tibialis* and *P. confusa*, the two most abundant parasitoid species, were widespread all along the latitudinal gradient while *S. bella* was absent from the Stockholm area (Table 1, Fig. 2). We also found *P. vulgaris* and *M. subcompleta* in most of the sampling sites and across the three counties (Table 1, Fig. 2). We recorded other parasitoid species in low numbers, which, therefore, provide limited information about their latitudinal distribution. Still *T. haereticus* (n=21) was restricted to the two northern counties and *C. vanessae* (n=30) to the two southern counties (Table 1, Fig. 2).

**Table 1.** Showing the distribution of larvae dead, according to sampling sites and counties, by parasitoid family and species, and due to unknown causes, covering infection by virus, bacteria, or fungi. The sites are ordered latitudinally. Note that 5 larvae were parasitized by two different species, which lead to the discrepancy between the total by family and the grand total.

| Larval death by: |  | Tachinids |  |  |  |  |  | Ichneumonids |  |  | Braconids |  | parasitoid<br>not<br>identified | unknown<br>causes | Grand<br>Total |
| --- | --- | --- | --- | --- | --- | --- | --- | --- | --- | --- | --- | --- | --- | --- | --- |
| Species |  | *Pelatachina* | *Sturmia* | *Phryxe* | *Phryxe* | *Pales* | *Blondelia* | *Phobocampe* | *Thyrateles* | *Thyrateles* | *Microgaster* | *Cotesia* |  |  |  |
| County | Sites | *tibialis* | *bella* | *vulgaris* | *nemea* | *pavida* | *nigripes* | *confusa* | *haereticus* | *camelinus* | *subcompleta* | *vanessae* |  |  |  |
| Stockholm | 235 | 29 |  | 5 |  |  |  | 18 | 6 |  | 1 |  |  | 31 | 90 |
|  | 284 | 40 |  | 4 |  |  |  | 10 |  |  | 9 |  | 3 | 40 | 106 |
|  | 569 | 29 |  | 1 |  |  |  | 20 | 5 | 1 | 3 |  | 6 | 132 | 197 |
|  | 631 | 10 |  | 2 |  |  |  | 62 | 1 |  | 2 |  | 7 | 127 | 211 |
|  | 390 | 14 |  | 13 |  |  |  | 24 | 5 |  | 2 |  | 4 | 65 | 127 |
|  | 590 | 18 |  | 1 |  |  |  | 30 | 1 |  | 7 |  | 7 | 42 | 106 |
| Kronoberg | 31 |  | 5 | 1 |  |  |  |  |  |  |  |  | 5 | 8 | 19 |
|  | Ljungby | 23 | 6 |  |  |  |  | 3 |  |  | 2 |  | 3 | 24 | 61 |
|  | Odensjö | 6 | 19 |  |  | 1 |  |  |  |  | 1 |  |  | 6 | 33 |
|  | 37 | 24 | 18 | 3 |  |  | 4 | 26 |  |  | 1 | 1 | 15 | 57 | 148 |
|  | 178 | 18 | 3 | 4 |  |  |  | 40 | 3 |  | 2 |  | 4 | 35 | 109 |
| Skåne | 19 | 14 | 2 | 3 | 2 |  |  | 4 |  |  | 6 |  | 1 | 2 | 34 |
|  | Åsvägen | 5 | 14 |  |  |  |  | 1 |  |  | 9 |  |  | 13 | 42 |
|  | 662 | 73 | 17 | 8 |  |  | 1 | 14 |  |  | 2 | 3 | 2 | 24 | 142 |
|  | 915 | 16 | 17 | 1 |  |  |  | 5 |  |  | 16 | 1 | 3 | 27 | 86 |
|  | 63 | 32 |  |  |  |  |  | 18 |  |  | 13 |  | 5 | 16 | 84 |
|  | 335 | 64 | 43 | 9 |  |  |  | 33 |  |  | 17 | 4 | 16 | 42 | 227 |
|  | 947 | 23 | 51 |  |  |  |  | 21 |  |  | 7 |  | 25 | 49 | 176 |
|  | 6 | 88 | 10 | 4 | 1 |  |  | 101 |  |  | 14 | 21 | 12 | 60 | 310 |
| Subtotal |  | 526 | 205 | 59 | 3 | 1 | 5 | 430 | 21 | 1 | 114 | 30 | 118 | 799 |  |
| Total |  | 799 |  |  |  |  |  | 452 |  |  | 144 |  | 118 | 799 | 2308 |

**Table 2.** Table showing the numbers of butterfly larvae of each species dead according to parasitoid species or due to unknown causes, which cover infection by virus, bacteria, or fungi. The table also summarizes the contribution of each parasitoid species to the total parasitism found per butterfly species and intermediate summaries show parasitoids contribution by family. The percentages of larvae dead due to unknown causes are related to the total amount of larvae of each sampled species.

| Host butterfly / parasitoid species | *A. urticae*<br>(n) | *A. io*<br>(n) | *V. atalanta*<br>(n) | *A. levana*<br>(n) | **Total**<br>**(n)** | *A. urticae*<br>(%) | *A. io*<br>(%) | *V. atalanta*<br>(%) | *A. levana*<br>(%) |
| --- | --- | --- | --- | --- | --- | --- | --- | --- | --- |
| *Pelatachina tibialis* | 312 | 207 | 7 |  | **526** | 45.4 | 34.5 | 4.3 |  |
| *Sturmia bella* | 39 | 105 | 10 | 51 | **205** | 5.7 | 17.5 | 6.1 | 82.3 |
| *Phryxe vulgaris* | 32 | 18 | 9 |  | **59** | 4.7 | 3.0 | 5.5 |  |
| *Blondelia nigripes* |  | 5 |  |  | **5** |  | 0.8 |  |  |
| *Phryxe nemea* | 1 |  | 2 |  | **3** | 0.1 |  | 1.2 |  |
| *Pales pavida* |  |  | 1 |  | **1** |  |  | 0.6 |  |
| **Total tachinids** | **383** | **335** | **29** | **51** |  | **55.9** | **55.8** | **17.7** | **82.3** |
| *Phobocampe confusa* | 229 | 197 |  | 2 | **428** | 33.3 | 32.8 | 0.0 | 3.2 |
| *Thyrateles haereticus* | 8 | 11 | 2 |  | **21** | 1.2 | 1.8 | 1.2 |  |
| *Thyrateles camelinus* |  |  | 1 |  | **1** |  |  | 0.6 |  |
| *Campopleginae, Diadegma sp* | 1 |  | 1 |  | **2** | 0.1 |  | 0.6 |  |
| **Total ichneumonids** | **238** | **208** | **4** | **2** |  | **34.6** | **34.7** | **2.4** | **3.2** |
| *Microgaster subcompleta* | 1 |  | 113 |  | **114** | 0.1 |  | 68.9 |  |
| *Cotesia vanessae* | 24 |  | 6 |  | **30** | 3.5 |  | 3.7 |  |
| **Total braconids** | **25** | **0** | **119** | **0** |  | **3.6** | **0.0** | **72.6** | **0.0** |
| **Unknown parasitoids** | **40** | **57** | **12** | **9** | **118** | **5.8** | **9.5** | **7.3** | **14.5** |
| **Unknown causes** | **205** | **447** | **83** | **64** | **799** | **9.1** | **19.8** | **12.2** | **4.2** |

**Figure 2.**
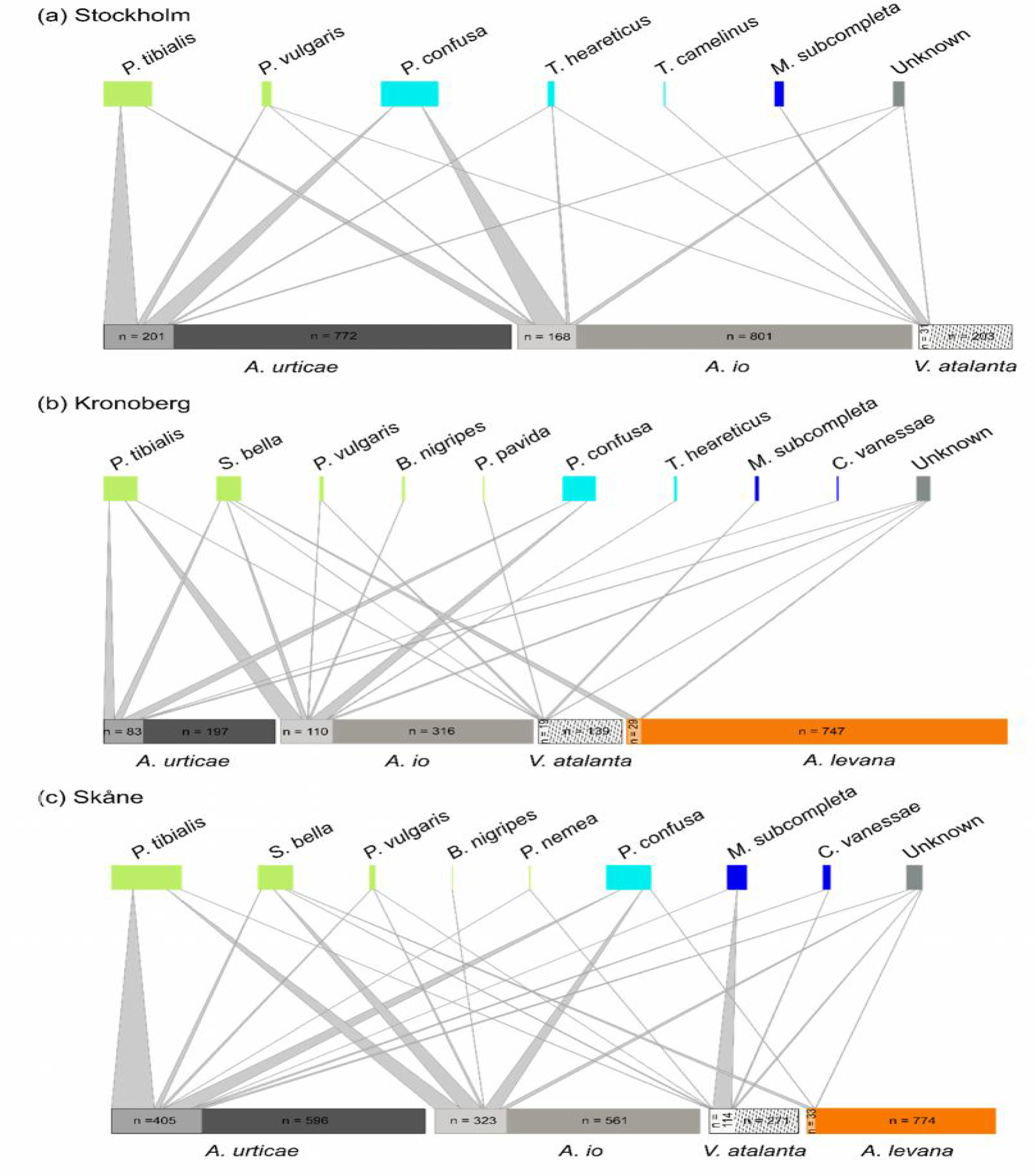
Quantitative host butterflies-parasitoid species association in the counties of (a) Stockholm, (b) Kronoberg, (c) Skåne. For each web, the bottom boxes represent, for each butterfly species, the proportion of larvae parasitized within the total amount of larvae sampled, the upper boxes correspond to the contribution of each parasitoid species to the overall parasitism. Associations are ordered according to parasitoid family.

The parasitoid complex varied among the butterfly hosts. *Vanessa atalanta* was the host of most parasitoid species including representatives of all three families (Table 2). *Aglais urticae* was also found to be parasitized by a wide range of species from the three families (Table 2). *Aglais io* and *A. levana* were not parasitized by braconids and *A. levana* was almost exclusively parasitized by *S. bella*, except on two occasions by *P. confusa* (Table 2). Note that the three most abundant parasitoid species were shared among the butterfly hosts, except for *P. tibialis* and *P. confusa* that were never observed in *A. levana* and *V. atalanta* larvae, respectively. We also recorded cases where the cause of larval death was unknown. While, to a certain extent, we relate this mortality to parasitoids that failed to achieve their development within the body of their host either due to a late attack of the parasitoid or to the immune response of their host (HA, personal observation), we also recorded cases of mortality due to viral infection, bacteria or fungi. The overall percentage of dead larvae due to unknown causes varied from 4.2% for *A. levana* to 19.8% for *A. io* (Table 2). The high mortality of *A. io* is not surprising as this species is relatively sensitive to laboratory rearing conditions, especially during the early instars (HA, personal observation).

### Effects of latitude and phenology on parasitism rates

Parasitism was responsible for high mortality, particularly in *A. urticae* and *A. io* (Fig. 3a, Table S4) and showed a gradual decrease along the latitudinal gradient, from Skåne to Stockhom (Fig. 3a). Over the two field seasons, 40.2% of *A. urticae* and and 37.0% of *A. io* larvae collected in Skåne were parasitized. These rates decrease to 20.4% and 17.4% in Stockholm County for *A. urticae* and *A. io*, respectively. *Aglais urticae* showed higher parasitism rates than *A. io*, although this effect is driven mainly by the difference observed in the Stockholm area (Fig. 3a, Table S4). Across counties, *A. urticae* and *A. io* were parasitized at significantly higher frequency than *V. atalanta* and *A. levana* (Fig. 3a). Over the two field campaigns, *V. atalanta* showed highest parasitism rate in Skåne, with 39.9% of the larvae collected parasitized, while it was 12.0% and 13.1% in the counties of Kronoberg and Stockholm, respectively. *Araschnia levana* was very weakly parasitized, with parasitism rates of 4.1% in Skåne and 3.9% in Kronoberg.

**Figure 3.**
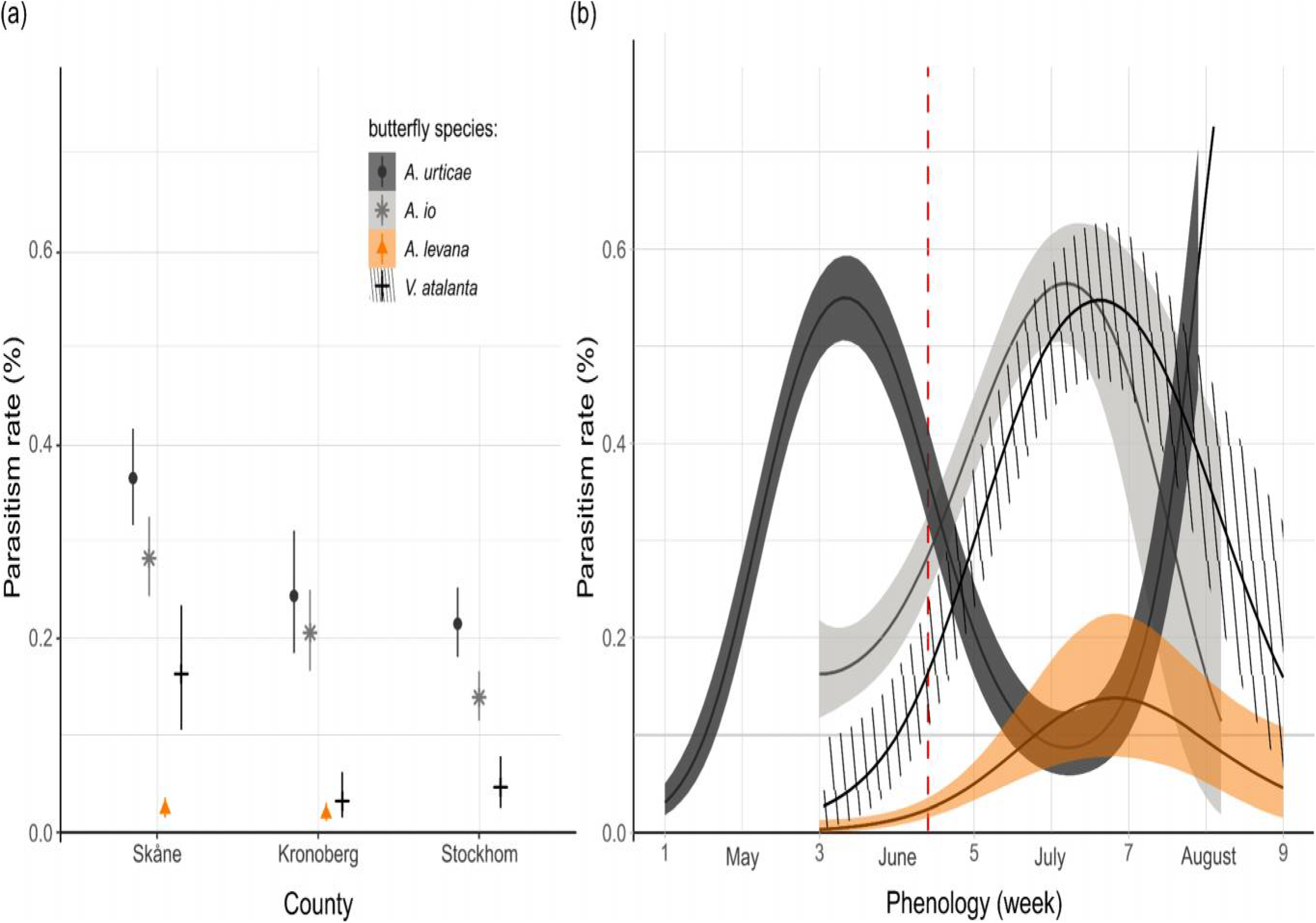
(a) Estimation of marginal means of parasitism rates (%) at representative values (week = 4.4, year = 2017) according to butterfly species and counties (mean and 95% confidence interval). (b) Estimated variation in the parasitism rate by species over time (weeks) in 2017 in Skåne. Non-overlapping confidence intervals correspond to significant differences in parasitism rate between groups. Note that we have adjusted for week 4.4 as at this time, differences in parasitism between species reflect the overall differences observed the season. The phenology of parasitism is illustrated in Skåne but follows the same pattern in the other two counties, modulated by a variation in the intercept. The red line on (b) indicates week 4.4, the time of the reproductive season for which the marginal means shown in (a) were extracted for Skåne. We restricted the plot of estimated variation in parasitism rate to the time window for which each species was sampled in the field.

While the overall parasitism rate was significantly lower in 2017 compared to 2018 (estimate = −0.33, 95% CI = [−0.17, −0.49], Table S4), within each season, parasitism was also lower in early batches than in the later ones. The seasonality of parasitism was, however, specific to each butterfly species (Fig 3b, Table S4) and results from differences in their phenology and the phenology of their parasitoids. Parasitism rate in *A. urticae* followed a bimodal distribution that reflects the bivoltine life cycle of the species in Sweden. In contrast, parasitism rate in *A. io* and *V. atalanta* followed a unimodal pattern with a peak at the end of July. We observed a similar unimodal pattern of parasitism in *A. levana* but the low parasitism in this species makes it difficult to form reliable estimates of its phenological variations.

### Effect of butterfly species assemblage on parasitism rates

The impact of community composition, that is, the number and identity of co-occurring larval species and the total abundance of larvae, on rates of parasitism is specific to each species (Fig. 4, Table S5).

**Figure 4.**
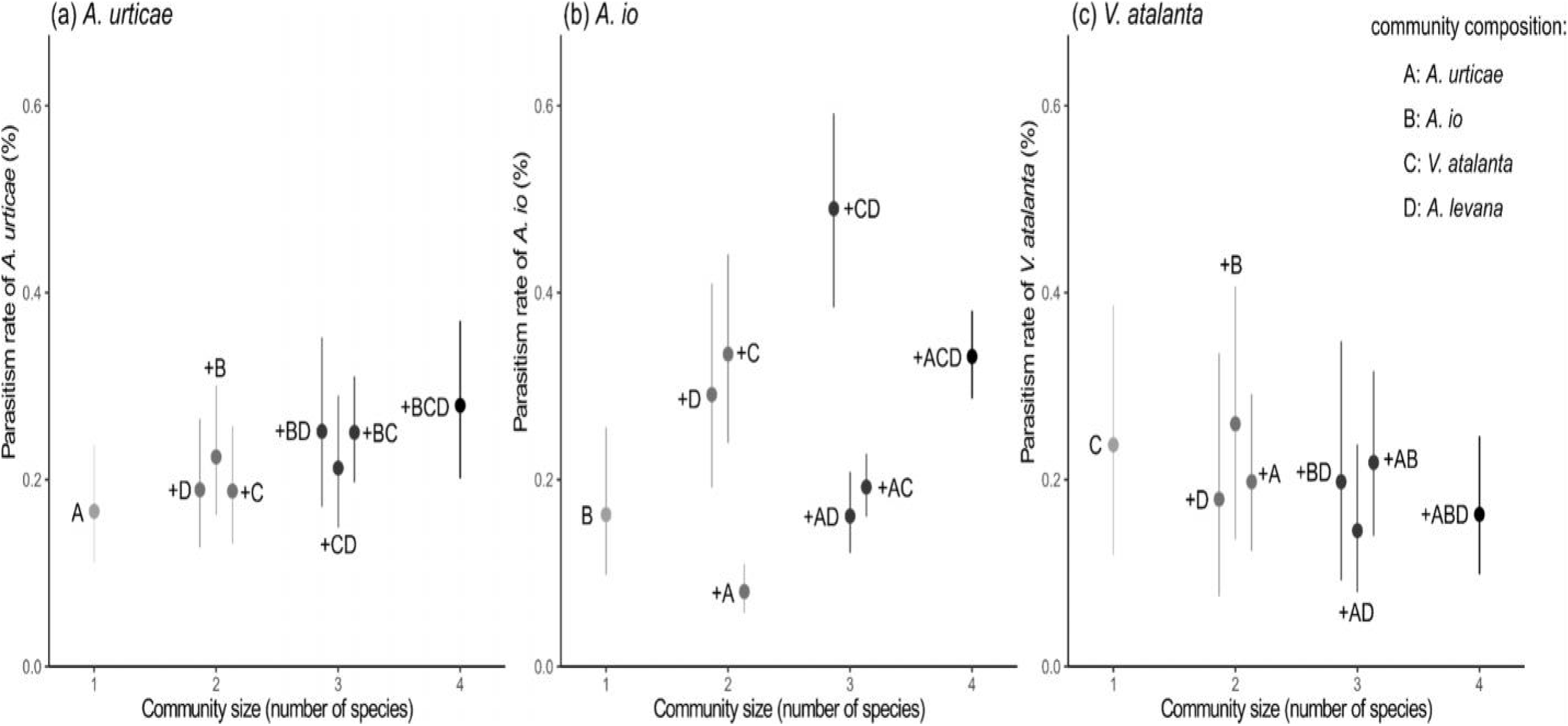
Contrasting effects of community composition, taken as the presence/absence of the other species, including *A. levana*, on parasitism rate of (a) *A. urticae*, (b) *A. io*, and (c) *V. atalanta*. Estimation of marginal means of parasitism rates (%) are given at representative values (week = 4.74) and parasitism rates of each of the focal species are ordered on the x-axis according to the number of species which co-occur. The first bar on each plot corresponds to parasitism rate of the focal species found alone (mean ± CI) at each site and the letter stands for the identity of the focal species with A for *A. urticae*, B for *A. io*, C for *V. atalanta*. The following bars correspond to parasitism rate of the focal species (mean ± CI) when co-occurring with other nettle-feeding butterflies with +A when the species co-occur with *A. urticae*, +B with *A. io*, +C with *V. atalanta*, and +D with *A. levana*. Non overlapping confidence intervals correspond to significant differences in parasitism rate between groups.

Parasitism in *A. urticae* is higher when larvae are abundant (estimate = 0.26, 95% CI = [0.09, 0.42], Fig. 4a, Table S5) and is elevated when *A. urticae* co-occurs with *A. io* (estimate = 0.40, 95% CI = [0.05, 0.76], Fig. 4a). Parasitism in *A. io* was not sensitive to the abundance of larvae at the time of collection (estimate = −0.03, 95% CI = [−0.17, 0.10], Fig. 4b, Table S5) but varied according to species assemblage and community size (Fig. 4b). In particular, parasitism rate in *A. io* increased when co-occurring with *V. atalanta* (estimate = 1.05, 95% CI = [0.73, 1.38]) and *A. levana* (estimate = 0.82, 95% CI = [0.57, 1.07]), and decreased when co-occurring with *A. urticae* (estimate = −0.83, 95% CI = [−1.32, −0.34], Table S5). We also observed that parasitism rate in *A. io* increased with the number of co-occurring species (Fig. 4b). We did not observed an effect of larvae abundance or species assemblage on parasitism in *V. atalanta* (Fig. 4c, Table S5).

### Parasitism rate and time since establishment of A. levana

The time period since first observation of *A. levana* significantly explained variations in parasitism rate in *A. urticae* and *A. io* (LR Chisq(3) = 35.15, p < 0.001 for *A. urticae* and LR Chisq(3) = 15.88, p = 0.001 for *A. io*, Fig. 5, Appendix S5), which showed higher parasitism rates in the earliest colonized sites. Parasitism in *A. io* additionally decreased with latitudinal (LR Chisq(1) = 5.22, p = 0.022, Appendix S5). Parasitism in *V. atalanta* was not explained by differences in the time period since first observation of *A. levana* but decreased with latitude (LR Chisq(1) = 24.61, p < 0.001, Appendix S5).

**Figure 5.**
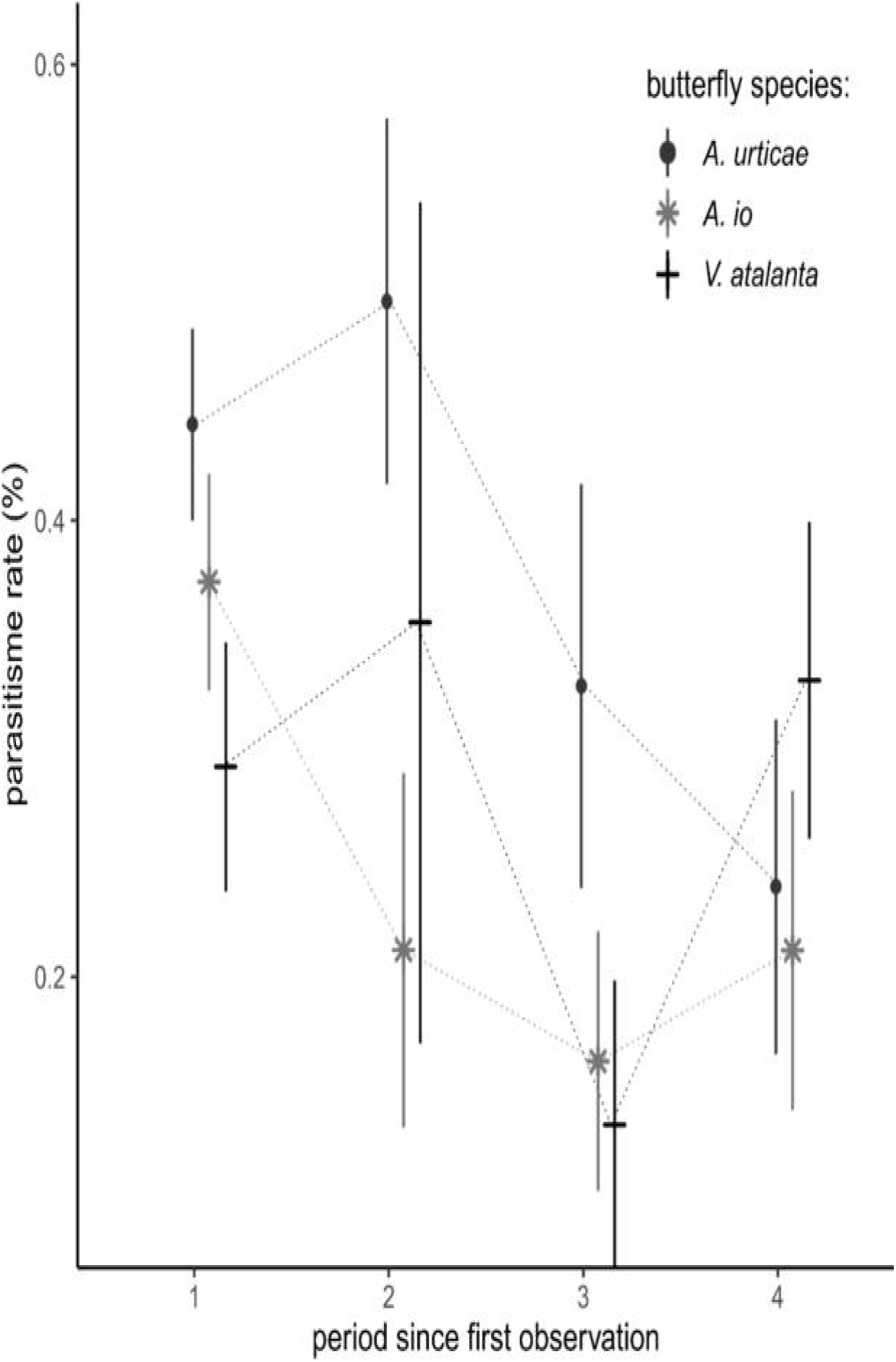
Parasitism rate (mean ± se) of *A. urticae, A. io*, and *V. atalanta*, according to the time period of establishment of *A. levana* at the site. The four time periods correspond to the division of the distribution of the time since first observation of *A. levana* into four quantiles and are ordered chronologically.

## Discussion

Our results highlight the influence of species assemblages and trophic interactions on the parasitism of nettle-feeding butterflies. We showed that parasitism was responsible for high mortality rates in two of the native species, *A. urticae* and *A. io*. In comparison, parasitism caused lower mortality in *V. atalanta* and *A. levana*. The parasitoid complex was shared among the nettle-feeding butterflies but *A. levana*, the newcomer in Sweden, was almost exclusively parasitized by the tachinid *S. bella*. We observed that parasitism was influenced by community composition and that this effect was specific to each butterfly species. In addition, we found higher rates of parasitism in the native species at sites where *A. levana* has established for a longer time period.

The low parasitism rate in *V. atalanta* and *A. levana* might be the result of morphological, physiological, behavioural, and immunological differences, compared to the other study species. *Vanessa atalanta* larvae are solitary, which may complicate the search for host larvae by their parasitoids, in comparison to the other species that lay batches of eggs (Gentry & Dyer, 2002; Hawkins, 2005). However, *V. atalanta* larvae also live concealed in folded leaves, a shelter-building behaviour that has been shown to concentrate chemical and visual signals that facilitate the localization of individual larvae by parasitoids (Dyer & Gentry 1999). Alternatively, differences in feeding guilds between butterflies (solitary versus gregarious) may influence preference of host of parasitoids, as the result of the evolution of different strategies of host searching behaviours in parasitoids, and participate to explain the lower attack rate in *V. atalanta* (Sigiura, 2007). Nevertheless, the low parasitism measured in *V. atalanta* is difficult to explain, considering that this species was host for the largest diversity of parasitoids and that it has been documented to be highly parasitized in other parts of its range (see Rice, 2012). Variation in *V. atalanta* parasitim rates across its range might be related to its migratory behaviour, conditions at overwintering sites, and synchrony between the butterfly and its parasitoids. The pattern is different for *A. levana*, which is resident in Sweden and has been found to be weakly parasitized in other parts of its distribution (Wagner et al., 2011). *Araschnia. levana* larva show a pronounced dropping behaviour, which in other species has been shown to be effective against parasitoids which lose track of the chemical and sensory cue of their hosts (Gross and Price, 1988; Fitzpatrick et al., 1994). Alternatively, lower parasitism in *A. levana* could be a result of its recent establishment in Sweden. The enemy release hypothesis (Jeffries & Lawton, 1984; Keane & Crawley, 2002) predicts that in a new area, species experience a period when they escape their natural enemies, until interactions with the local parasitic complex are established (Menéndez et al., 2008). In Sweden, *A. levana* was first reported in the 1980s but probably became established more recently, as there are very few reports of the species before 2000 (see Appendix S1). Considering the relatively short time that was available for recruitment of local parasitoids (Cornell & Hawkins 1992), we can not rule out the possibility that lower level of parasitism observed in *A. levana* are partly a consequence of its recent establishment and that populations have escaped from parasitism during its expansion phase. This hypothesis is strengthened by the fact that Söderlind (2009) reported no parasitism in *A. levana* in South Sweden, while our data reveal that the species has now been colonized by local parasitoid populations. Future monitoring of parasitism load in *A. levana* populations in Sweden and across the wider distribution range of the species would be necessary to disentangle the relative importance of these two hypotheses.

Butterfly community composition was significantly associated with parasitism in *A. urticae* and *A. io* but not in *V. atalanta*. The differences in egg-laying behaviour mentioned above, where parasitoids prefer gregarious species when present, is again one potential explanation, but *V. atalanta* was also mostly parasitized by *M. subcompleta*, a parasitoid associated almost exclusively with this species. *Aglais io* seemed to benefit from the co-occurrence of *A. urticae*, which was associated with reduced parasitism, while parasitism in *A. urticae* increased when it co-occurred with *A. io*. In contrast, parasitism in *A. io* is increased when co-occurring with*A. levana* and *V. atalanta*. Previous work on this study system hypothesized that a change in the composition of this community, namely, the arrival of *A. levana*, would influence the dynamics and spatial distribution of the resident butterflies through apparent competition (Audusseau et al., 2017). The association between host community composition and parasitism of *A. io* and *A. urticae* is consistent with this prediction. Parasitism rate of the native species decreased along the south-north gradient and was lower in sites recently colonized by *A. levana*, highlighting the potential role of *A. levana* in explaining the high parasitism rates of the native species in the southern counties. Nonetheless, while we found an increase in parasitism in *A. io* when the species co-occured with *A. levana*, this was not observed in *A. urticae* (Fig. 4a & b). The more pronounced shift in distribution of *A. urticae* reported by Audusseau et al. (2017) could have, however, suggested a relativly stronger response of parasitism in *A. urticae* to the co-occurrence of *A. levana*. The additional analysis that we propose in the Appendix S3 suggests, moreover, that parasitism in *A. urticae*, when restricted to parasitism caused by parasitoids shared with *A. levana*, is elevated when the species co-occurs with *A. levana*. Furthermore, parasitism in *A. urticae* increases with the total abundance of larvae, a phenomenon that might partly be associated with the arrival of the novel host. Differences in the phenology of parasitism between hosts also suggest that *A. levana* could provide a refuge for parasitoids at a time when the native hosts (*A. urticae* and *A. io*) are rare. Thus, species co-occurrence at a site over the season, rather than at a sampling event, may influence their level of parasitism. Last, our study focused on larval parasitoids (for reasons previously mentioned), but pupal parasitoids are known to be shared among our study butterflies and to cause high mortality (Pyörnilä, 1977; Shaw et al., 2009). In particular, the restricted host range of the pteromalid *Pteromalus puparum*, which includes the butterflies of our study (Shaw et al., 2009), and the size of its brood, make this species a strong candidate for driving apparent competition in our study community.

From our study, we can not rule-out the effect of other differences across counties, such as changes in parasitoid species richness, population dynamics, habitat quality, or variation in phenological synchrony between the butterflies and their parasitoids, which may all contribute to explain the latitudinal decrease in parasitism. For example, the occurrence of other hosts over the landscape may influence the population dynamics of parasitoids and mediate apparent competition (Davis, 1991; Gaston, 2005). Parasitoids are also responding to the conditions of their habitat (Shaw 2006), which may vary between counties, despite our effort to select sites with comparable landscape. The latitudinal decrease in parasitism could also be associated with a latitudinal trend in weather conditions. Temperature affects insect-parasitoid interactions (Thomas & Blandford, 2003). While in some systems parasitoid activity can increase with temperature (Mann et al., 1990), which could lead to a higher activity period and oviposition rate in the parasitoids at lower and warmer latitude, the literature does not provide consistent evidence of such a pattern (Hawkins, 2005). In our system, the differences in microclimatic conditions across sites did not align with the latitudinal pattern observed for parasitism (see Appendix S7), but we observed latitudinal differences in the parasitoid community. For example, *S. bella*, one of the most abundant parasitoid species in our sample, was only found in the two southern counties. It has also recently established in the U.K. and its arrival coincided with the decline of *A. urticae*. However, Gripenberg et al. (2011) were not able to provide clear support for the role of *S. bella* in the decline of *A. urticae*. Manipulative experiments on community composition while controlling for host abundances would shed further light on parasitoid host preferences and on the mechanism of apparent competition, such as how the parasitoid population built up throughout the season.

The systematic sampling that we carried out in the field, at these temporal and spatial scales, and on a set of species that are assembled in a community is rare, but crucial to further our understanding of indirect biotic interactions that structure the community and their persistence and stability over time and space. It enabled us to study the manner by which species composition, variation in abundance, species phenology, and the arrival of *A. levana*, influence local biotic interactions and, ultimately, provide evidence consistent with the role of apparent competition mediated by shared parasitoids in nettle-feeding butterflies. In particular, we showed that parasitoid pressure plays a major role, having an important effect on mortality of our study species in Sweden. We also provide further evidence that modifications favourable to the population dynamics of parasitoids, such as the arrival and establishment of *A. levana*, has the potential to modify the pressure parasitoids exert on their native hosts. Hence, modification of the biotic interactions should be further studied to assess the full impact of environmental change on populations and communities. As mentioned elsewhere (Gaston, 2010), this is all the more important in common species as their ubiquity and abundance often makes them connect with a large number of species through trophic interactions.

## Acknowledgments

H. Audusseau acknowledges support from the Swedish Research Council (2016-06737). This work benefited from technical assistance by Maria Celorio-Mancera de la Paz and Houshuai Wang, from taxonomic assistance on the tachinids by Christer Bergström, and from genetic assistance (for molecular confirmation of a subset of the parasitoids determined) by Lise Dupont and Claire Brice. Tom August provided the script for the *A. levana* range expansion animation. We are also grateful to all reporters of the Swedish Biomonitoring Scheme and specifically to Evald Jonsson, Sven Nilsson, Peter Rolfson, Pål Axel Olsson, Kurt Norell, and Mats Hansson for valuable advice on the distribution of species at the sites.

## Data Accessibility

Data presented in this manuscript will be available upon acceptance.

## Supporting information

### Appendix S1

Animation showing the expansion of *Araschnia levana* over the period 1995-2018. The data used are the opportunistic occurrence data collected by amateurs and available from Artportalen (Swedish Species Observations System, www.artportalen.se). The black dots correspond to species occurrences, red dot correspond to new occurrences as time goes. The red halo around each new observation is expanding until it vanishes after 200 days.

### Appendix S2 Details on the winter diapause conditions

In mid-September, we stored the moist plastic vials containing the pupae of *A. levana* and parasitoids at +8 °C. In November, we transferred them to a climate chamber with a day/night temperature of +4 / 0°C and a light regime of 12L:12D and changed the day/night temperature to −4 / −2 ° C from mid-December to the end of February. Subsequently, we reversed the temperature cycle by following the same temperature scheme. Throughout the hibernation period, we frequently checked the moisture conditions and adjusted them if necessary. We broke diapause of the pupae of *A. levana* and of the parasitoids around mid-April. To do this, we transferred the plastic vials to ambient temperature and light conditions, and sprayed them with water regularly so that the individuals rehydrated.

### Appendix S3 Effect of butterfly species assemblage on each of the native species’ parasitism by the subset of shared parasitoid species with *A. levana*

#### Aim and dataset

Apparent competition among species can only occur when species share a natural enemy, that is, parasitoids in the case of our butterfly community. From our field sampling, while most parasitoid species attack different host butterflies, the only parasitoid that is shared by all four study butterflies is *S. bella* (Table 2), and would, therefore, be the only one species potentially involved in apparent competition between our study butterflies.

**As our aim is to provide further evidence consistent with the fact that the establishment of** *A. levana* **may have increased parasitism in the native species, as a result of apparent competition, we provide, here, an extra analysis on parasitism by shared parasitoids between the native species and** *A. levana*. Restricting the analysis of the effect of species assemblage on parasitism by *S. bella* would, however, drastically reduce the dataset to use. First, because *S. bella* is only found in the two southern counties^1^. Second, this could also lead, for consistency, to restrict our analysis to the phenological window of occurrence of the *S. bella* (five sampling occasions per site in 2017).

Considering that other parasitoids have been documented as attacking *A. levana* (they are *Apechthis compunctor*, *Thyrateles camelinus*, *Compsilura concinnate*, *Phryxe nemea*, *Phryxe vulgaris*, and *Sturmia bella*, Herting & Simmonds, 1976), **we will examine the effect of butterfly assemblage on parasitism by the following subset of parasitoids, *Thyrateles camelinus*, *Phryxe nemea*, *Phryxe vulgaris*, *Sturmia bella***, and** *Phobocampe confusa***, on the basis that these parasitoids were also collected over our study sites. These analyses are limited to the two southern counties (Sk*åne and Kronoberg)*, where *A. levana* occurs.

#### Analyses

These analyses followed the same procedure as described in the main text. We tested for the effect of the presence or absence of each species of butterfly, taken as a binary variable (0/1), and the effect of the abundance of larvae, on the parasitism rate of each of the native species. We also included in each model the non-linear effect of the sampling week (with k up to 4), to capture phenological variations of parasitism of each species. The abundance of larvae corresponds to the total number of larvae from all species collected per site and sampling week and was zero-centred prior to inclusion in the models. We performed these analyses in a Bayesian framework using generalized linear and nonlinear multivariate multilevel models. Parasitism was modelled assuming a zero inflated binomial distribution with a logit link function. The models were fitted through MCMC sampling, using the Hamiltonian Monte Carlo algorithm implemented in Stan (Carpenter et al., 2017) and the R interface provided in the brms package (Bürkner 2017; 2018). We ran four chains for 10000 iterations with the first 4000 discarded as burn-in and used the default priors. To test for significant differences in parasitism between county and species, we compared the posterior probability distribution of the model parameters.

#### Results

We observed that the number and identity of co-occurring larval species and the total abundance of larvae affected parasitism by the subset of parasitoids considered in this analysis, and was specific to each species (Fig. S3, Table S3). However, the patterns of variation of each species’ parasitism rates are different from the observed patterns when including all parasitoid species and the three counties.

Parasitism in *A. urticae* is higher when the species co-occurs with *A. levana* (estimate = 1.01, 95% CI = [0.31, 1.70], Fig. S3, Table S3). Parasitism in *A. io* no longer showed sensitivity to the co-occurrence of *A. urticae* and *A. levana*, but still increased when the species co-occurs with *V. atalanta* (estimate = 3.46, 95% CI = [2.51, 4.48], Fig. S3, Table S3). Parasitism in *A. io* is also higher when larvae are abundant (estimate = 0.65, 95% CI = [0.35, 0.96]). Similarly to the analysis present in the main manuscript, we did not observed an effect of larvae abundance or species assemblage on parasitism in *V. atalanta* (Fig. S3, Table S3).

**This analysis showed that, when focusing on the specific subset of parasitoids known to be shared between the native species and** *A. levana***, parasitism in** *A. urticae* **is increased when the species co-occurs with** *A. levana*.

---

**Figure S3.**
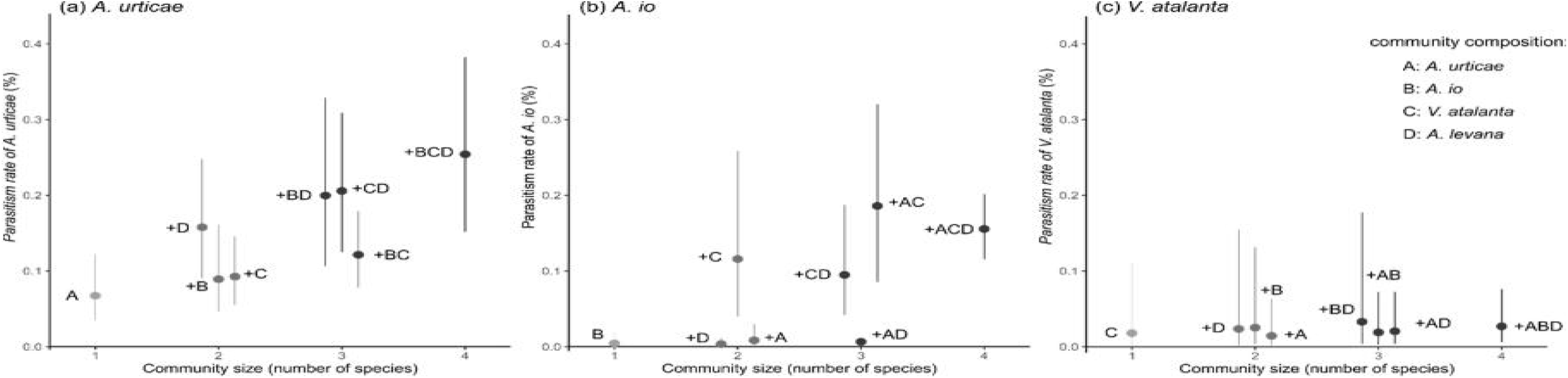
Contrasting effects of community composition, taken as the presence/absence of the other species, including *A. levana*, on parasitism rate by of (a) *A. urticae*, (b) *A. io*, and (c) *V. atalanta*. Only parasitism by the following parasitoid species is considered here: *Thyrateles camelinus*, *Phryxe nemea*, *Phryxe vulgaris*, *Sturmia bella*, and *Phobocampe confusa*. Estimation of marginal means of parasitism rates (%) are given at representative values (week = 4.74) and parasitism rates of each of the focal species are ordered on the x-axis according to the number of species which co-occur. The first bar on each plot corresponds to parasitism rate of the focal species found alone (mean ± CI) at each site and the letter stands for the identity of the focal species with A for *A. urticae*, B for *A. io*, C for *V. atalanta*. The following bars correspond to parasitism rate of the focal species (mean ± CI) when co-occurring with other nettle-feeding butterflies with +A when the species co-occur with *A. urticae*, +B with *A. io*, +C with *V. atalanta*, and +D with *A. levana*. Non overlapping confidence intervals correspond to significant differences in parasitism rate between groups.

**Table S3.** Summary table of the population-level effects of the presence/absence of each species of butterfly, the abundance of larvae, and seasonality (week of sampling), and of the non-linear effect of the seasonality (sds), on parasitism rates of the focal species. Estimates are provided on the logit-scale. Non overlapping confidence intervals correspond to significant differences in parasitism rate between groups. We assessed model fit by checking that the chains have mixed well and by looking at the distribution of the predictive values. Sds corresponds to the variance parameter (higher values reflecting more wiggly smoother). Note that the confidence intervals of the smooth terms are not overlapping zero. The smooth term is, therefore, required over the linear parametric effects of the week (see sweek: species). “zi” corresponds to the zero inflated estimate. The zero inflated binomial distribution model has two parts, a binomial count model and the logit model for predicting excess zeros.

| Butterfly species | variables | Estimate | 95% CI | Eff. Sample | Rhat |
| --- | --- | --- | --- | --- | --- |
| *A. urticae* | Intercept | -1.57 | [-1.87, -1.28] | 10223 | 1 |
|  | Presence *V. atalanta* | 0.36 | [-0.15, 0.87] | 10805 | 1 |
|  | Presence *A. io* | 0.32 | [-0.14, 0.81] | 10890 | 1 |
|  | Presence *A. levana* | 1.01 | [0.31, 1.70] | 11162 | 1 |
|  | Larvae abundance | 0.06 | [-0.27, 0.40] | 10412 | 1 |
|  | week | 1.90 | [1.14, 2.66] | 10495 | 1 |
|  | sds(sweek) | 16.32 | [6.81, 37.26] | 8118 | 1 |
|  | zi | 0.30 | [0.19, 0.41] | 10712 | 1 |
| *A. io* | Intercept | -4.33 | [-5.72, -2.99] | 8323 | 1 |
|  | Presence *V. atalanta* | 3.46 | [2.51, 4.48] | 7498 | 1 |
|  | Presence *A. urticae* | 0.63 | [-0.24, 1.53] | 10036 | 1 |
|  | Presence *A. levana* | -0.24 | [-1.05, 0.62] | 10089 | 1 |
|  | Larvae abundance | 0.65 | [0.35, 0.96] | 9070 | 1 |
|  | week | 2.24 | [-1.53, 7.56] | 4964 | 1 |
|  | sds(sweek) | 19.79 | [3.04, 52.66] | 6147 | 1 |
|  | zi | 0.41 | [0.30, 0.52] | 11010 | 1 |
| *V. atalanta* | Intercept | -3.35 | [-5.00, -1.84] | 10920 | 1 |
|  | Presence *A. io* | 0.38 | [-1.47, 2.29] | 10161 | 1 |
|  | Presence *A. urticae* | -0.23 | [-1.55, 1.13] | 10488 | 1 |
|  | Presence *A. levana* | 0.28 | [-0.99, 1.50] | 10335 | 1 |
|  | Larvae abundance | -0.06 | [-0.79, 0.64] | 9918 | 1 |
|  | week | 0.07 | [-2.56, 2.25] | 8209 | 1 |
|  | sds(sweek) | 7.54 | [0.36, 23.04] | 8582 | 1 |
|  | zi | 0.21 | [0.01, 0.56] | 10210 | 1 |

**Appendix S4.**
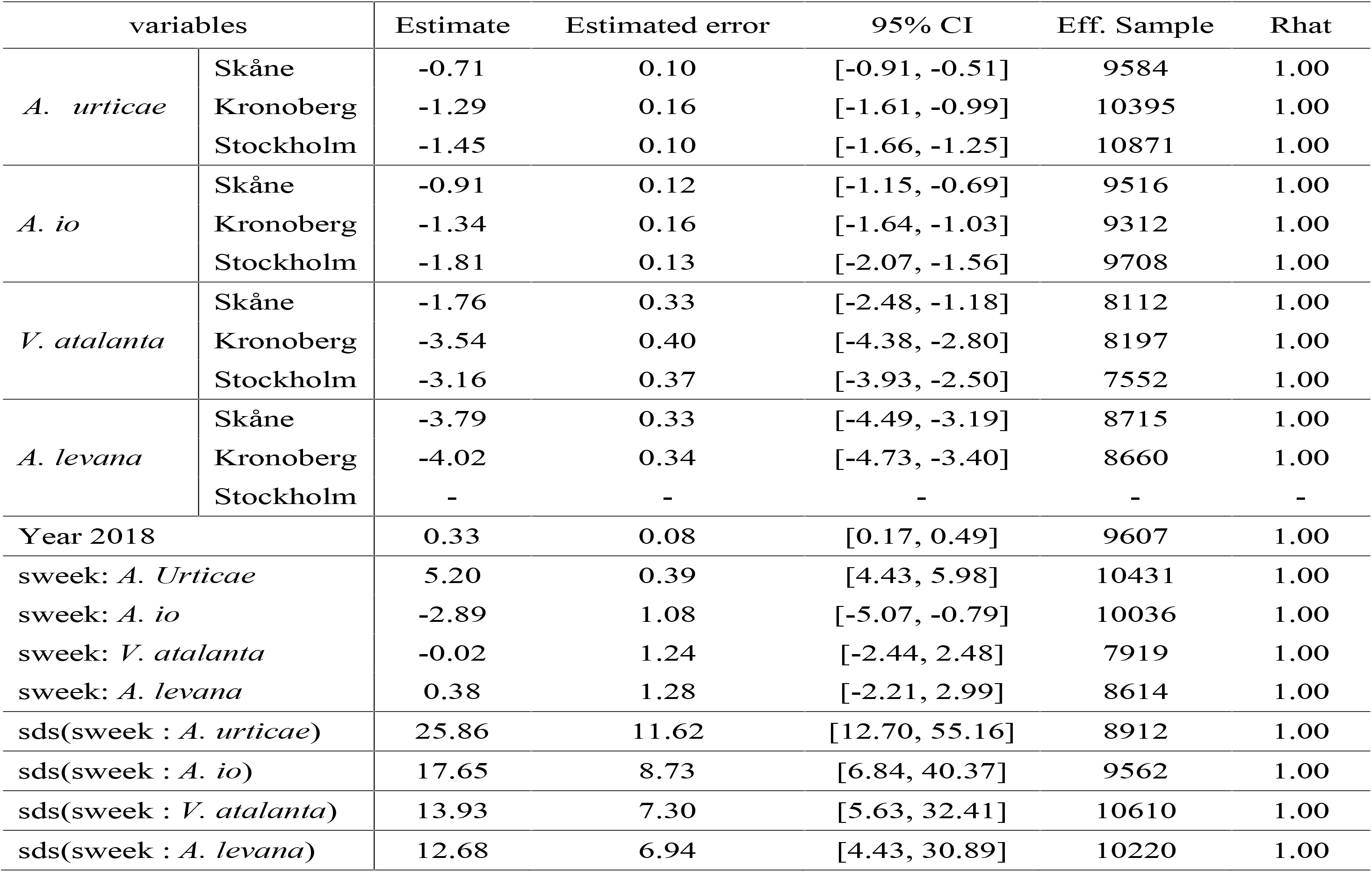
Summary table of the population-level effects of butterfly species, counties, sampling year, and seasonality (week of sampling), and of the non linear effect of the seasonality for each butterfly (sds), on parasitism rates. Estimates are provided on the logit-scale. Non overlapping confidence intervals correspond to significant differences in parasitism rate between groups. We assessed model fit by checking that the chains have mixed well and by looking at the distribution of the predictive values. Sds corresponds to the variance parameter (higher values reflecting more wiggly smoother). Note that the confidence intervals are not overlapping zero. The smooth term is, therefore, required over the linear parametric effects of the week (see sweek: species).

| variables |  | Estimate | Estimated error | 95% CI | Eff. Sample | Rhat |
| --- | --- | --- | --- | --- | --- | --- |
| *A. urticae* | Skåne | -0.71 | 0.10 | [-0.91, -0.51] | 9584 | 1.00 |
|  | Kronoberg | -1.29 | 0.16 | [-1.61, -0.99] | 10395 | 1.00 |
|  | Stockholm | -1.45 | 0.10 | [-1.66, -1.25] | 10871 | 1.00 |
| *A. io* | Skåne | -0.91 | 0.12 | [-1.15, -0.69] | 9516 | 1.00 |
|  | Kronoberg | -1.34 | 0.16 | [-1.64, -1.03] | 9312 | 1.00 |
|  | Stockholm | -1.81 | 0.13 | [-2.07, -1.56] | 9708 | 1.00 |
| *V. atalanta* | Skåne | -1.76 | 0.33 | [-2.48, -1.18] | 8112 | 1.00 |
|  | Kronoberg | -3.54 | 0.40 | [-4.38, -2.80] | 8197 | 1.00 |
|  | Stockholm | -3.16 | 0.37 | [-3.93, -2.50] | 7552 | 1.00 |
| *A. levana* | Skåne | -3.79 | 0.33 | [-4.49, -3.19] | 8715 | 1.00 |
|  | Kronoberg | -4.02 | 0.34 | [-4.73, -3.40] | 8660 | 1.00 |
|  | Stockholm | - | - | - | - | - |
| Year 2018 |  | 0.33 | 0.08 | [0.17, 0.49] | 9607 | 1.00 |
| sweek: *A. Urticae* |  | 5.20 | 0.39 | [4.43, 5.98] | 10431 | 1.00 |
| sweek: *A. io* |  | -2.89 | 1.08 | [-5.07, -0.79] | 10036 | 1.00 |
| sweek: *V. atalanta* |  | -0.02 | 1.24 | [-2.44, 2.48] | 7919 | 1.00 |
| sweek: *A. levana* |  | 0.38 | 1.28 | [-2.21, 2.99] | 8614 | 1.00 |
| sds(sweek : *A. urticae*) |  | 25.86 | 11.62 | [12.70, 55.16] | 8912 | 1.00 |
| sds(sweek : *A. io*) |  | 17.65 | 8.73 | [6.84, 40.37] | 9562 | 1.00 |
| sds(sweek : *V. atalanta*) |  | 13.93 | 7.30 | [5.63, 32.41] | 10610 | 1.00 |
| sds(sweek : *A. levana*) |  | 12.68 | 6.94 | [4.43, 30.89] | 10220 | 1.00 |

**Appendix S5.** Summary table of the population-level effects of the presence/absence of each species of butterfly, the abundance of larvae, and seasonality (week of sampling), and of the non linear effect of the seasonality (sds), on parasitism rates of the focal species. Estimates are provided on the logit-scale. Non-overlapping confidence intervals correspond to significant differences in parasitism rate between groups. We assessed model fit by checking that the chains have mixed well and by looking at the distribution of the predictive values. Sds corresponds to the variance parameter (higher values reflecting more wiggly smoother). Note that the confidence intervals are not overlapping zero. The smooth term is, therefore, required over the linear parametric effects of the week (see sweek: species). “zi” corresponds to the zero inflated estimate. The zero inflated binomial distribution model has two parts, a binomial count model and the logit model for predicting excess zeros. For example, the probability of *A. urticae* not being parasitized is actually higher that 0.23, but part of this probability is already modeled by the binomial distribution itself.

| Butterfly species | variables | Estimate | 95% CI | Eff. Sample | Rhat |
| --- | --- | --- | --- | --- | --- |
| *B. urticae* | Intercept | -0.66 | [-0.87, -0.46] | 10109 | 1 |
|  | Presence *V. atalanta* | 0.16 | [-0.18, 0.49] | 10615 | 1 |
|  | Presence *A. io* | 0.40 | [0.05, 0.76] | 9297 | 1 |
|  | Presence *A. levana* | 0.17 | [-0.30, 0.63] | 10970 | 1 |
|  | Larvae abundance | 0.26 | [0.09, 0.42] | 10154 | 1 |
|  | week | 3.55 | [2.95, 4.18] | 9662 | 1 |
|  | sds(sweek) | 24.54 | [11.86, 51.90] | 7052 | 1 |
|  | zi | 0.23 | [0.16, 0.30] | 10614 | 1 |
| *A. io* | Intercept | -1.17 | [-1.78, -0.56] | 11480 | 1 |
|  | Presence *V. atalanta* | 1.05 | [0.73, 1.38] | 11735 | 1 |
|  | Presence *A. urticae* | -0.83 | [-1.32, -0.34] | 11094 | 1 |
|  | Presence *A. levana* | 0.82 | [0.57, 1.07] | 11062 | 1 |
|  | Larvae abundance | -0.03 | [-0.17, 0.10] | 11308 | 1 |
|  | week | -0.04 | [-0.67, 0.55] | 8848 | 1 |
|  | sds(sweek) | 6.93 | [1.89, 18.95] | 7807 | 1 |
|  | zi | 0.22 | [0.15, 0.30] | 11479 | 1 |
| *V. atalanta* | Intercept | -0.85 | [-1.61, -0.07] | 11653 | 1 |
|  | Presence *A. io* | 0.15 | [-0.56, 0.89] | 11134 | 1 |
|  | Presence *A. urticae* | -0.27 | [-0.96, 0.40] | 11398 | 1 |
|  | Presence *A. levana* | -0.42 | [-1.05, 0.19] | 11082 | 1 |
|  | Larvae abundance | -0.09 | [-0.40, 0.21] | 10617 | 1 |
|  | week | 1.36 | [-0.05, 3.0] | 9383 | 1 |
|  | sds(sweek) | 10.74 | [3.83, 26.80] | 8409 | 1 |
|  | zi | 0.34 | [0.19, 0.47] | 10007 | 1 |

**Appendix S6.** Summary tables from the generalized linear models testing for each native butterfly species the effect of latitude and time since first observation of *A. levana*.

| Variables | *A. urticae* |  |  |  | *A. io* |  |  |  | *V. atalanta* |  |  |  |
| --- | --- | --- | --- | --- | --- | --- | --- | --- | --- | --- | --- | --- |
|  | Estimate | se | z | p-value | Estimate | se | z | p-value | Estimate | se | z | p-value |
| Intercept | -0.148 | 0.121 | -1.23 | 0.22 | -0.544 | 0.092 | -5.92 | **<0.001** | -1.083 | 0.195 | -5.56 | 0.14 |
| Latitude | 0.097 | 0.118 | 0.82 | 0.41 | -0.196 | 0.087 | -2.26 | **0.024** | -0.790 | 0.177 | -4.46 | **<0.001** |
| Period 2 | -0.155 | 0.182 | -0.85 | 0.40 | -0.775 | 0.212 | -3.66 | **<0.001** | -0.271 | 0.595 | -0.45 | 0.65 |
| Period 3 | -0.73 | 0.335 | -2.23 | **0.03** | -0.462 | 0.253 | -1.82 | 0.068 | 0.337 | 0.567 | 0.59 | 0.55 |
| Period 4 | -1.34 | 0.255 | -5.28 | **<0.001** | -0.114 | 0.216 | -0.53 | 0.60 | 0.610 | 0.281 | 2.17 | **0.03** |

**Appendix S7.**
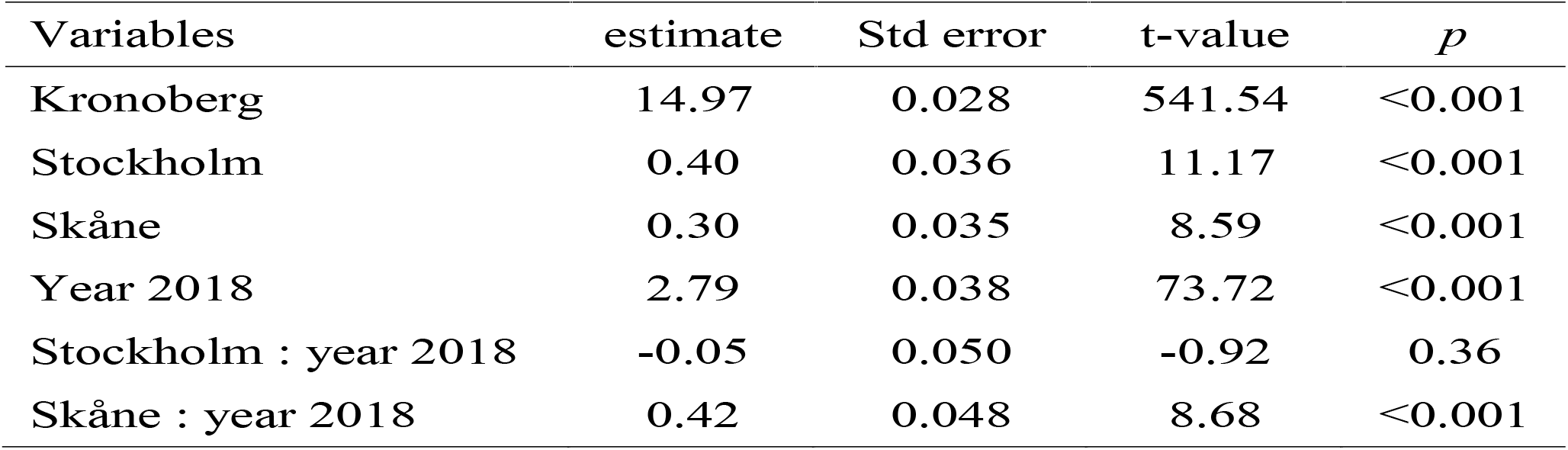
Summary table showing the effect of counties, years and the interaction between counties and years on the average temperature recorded by the temperature loggers placed in the field from May to August. We investigated differences in temperature averages between counties to test if these differences could explain the latitudinal pattern found in parasitism rates. We found that there is not a linear decrease in the average temperature as we go north. Indeed, the average temperature was +0.4°C higher in the Stockholm county than in Kronoberg. The the average temperature in 2018 is higher that in 2017, with on average +2.8°C. This increase was found to be slightly more pronounced in Skåne (estimate = 0.41, t-value = 8.7, p < 0.001).

| Variables | estimate | Std error | t-value | *p* |
| --- | --- | --- | --- | --- |
| Kronoberg | 14.97 | 0.028 | 541.54 | <0.001 |
| Stockholm | 0.40 | 0.036 | 11.17 | <0.001 |
| Skåne | 0.30 | 0.035 | 8.59 | <0.001 |
| Year 2018 | 2.79 | 0.038 | 73.72 | <0.001 |
| Stockholm : year 2018 | -0.05 | 0.050 | -0.92 | 0.36 |
| Skåne : year 2018 | 0.42 | 0.048 | 8.68 | <0.001 |

## Footnotes

1 Note on *S. bella*: the species has probably only recently established in Sweden from a different migration route than *A. levana*, as the first occurrence was reported on the East coast of Sweden (Christer Bergström, personal communication, with a first observation in Götland on 2003-07-24, coordinates: 57°31’52.7“N 18°08’52.4”E). Moreover, Söderlind (2009) reported no parasitism in *A. levana*, and therefore, no case of parasitism by *S. bella*.

